# Deep learning on reflectance confocal microscopy improves Raman spectral diagnosis of basal cell carcinoma

**DOI:** 10.1101/2022.03.03.482837

**Authors:** Mengkun Chen, Xu Feng, Matthew C. Fox, Jason S. Reichenberg, Fabiana C.P.S. Lopes, Katherine R. Sebastian, Mia K. Markey, James W. Tunnell

## Abstract

**Significance:** Raman spectroscopy may be useful to assist Mohs micrographic surgery for skin cancer diagnosis; however, the specificity of Raman spectroscopy is limited by the high spectral similarity between tumors and normal tissues structures such as epidermis and hair follicles. Reflectance confocal microscopy (RCM) can provide imaging guidance with morphological and cytological details similar to histology. Combining Raman spectroscopy with deep-learning-aided RCM has the potential to improve the diagnostic accuracy of Raman without requiring additional input from the clinician.

**Aim:** We seek to improve the specificity of Raman for basal cell carcinoma (BCC) by integrating information from RCM images using an Artificial Neural Network.

**Approach:** A Raman biophysical model was used in prior work to classify BCC tumors from surrounding normal tissue structures. 191 RCM images were collected from the same site as the Raman data and served as inputs to train two ResNet50 networks. The networks selected the hair structure images and epidermis images respectively within all the images corresponding to the positive predictions of the Raman Biophysical Model.

**Results:** Deep learning on RCM images removes 54% of false positive predictions from the Raman Biophysical Model result and keeps the sensitivity as 100%. The specificity was improved from 84.8% by using Raman spectra alone to 93.0% by integrating Raman spectra with RCM images

**Conclusions:** Combining Raman spectroscopy with deep-learning-aided RCM imaging is a promising tool to guide tumor resection surgery.

## 1 Introduction

Mohs micrographic surgery (MMS) is the most effective method to treat nonmelanoma skin cancer.^1,2^ With Mohs, the physician removes tumor tissue in stages. Within each stage, a layer of skin is excised and processed using frozen section histopathology, and the physician keeps removing layers until no tumor tissue is identified at surgical margins. Although Mohs has a cure rate greater than 98%,^2^ the histopathological analysis for each stage is relatively time-intensive (30-60min), requiring expensive clinical infrastructure and physician training. Optimization of care delivery using novel technology would add benefit for patients with high-risk skin cancer.

Raman spectroscopy (RS) is a promising tool for skin cancer diagnosis due to its high molecular specificity, low invasiveness, and low sample preparation requirement.^3^ RS is a sensing technique that uses light scattering to determine the molecular vibrational modes to provide chemical fingerprints.^4^ Prior studies have shown promising performance in differentiating Basal Cell Carcinoma (BCC) from surrounding normal tissues based on both analyzing the difference of BCC spectra from normal tissue spectra and using the nature and biochemical processes responsible for the spectral differences. Several recent studies report diagnostic sensitivities near 100% with specificities above 90%.^5–8^ Because Mohs is very successful with 98% cure rate, to be adopted as a low-cost alternative to Mohs, RS should also be highly accurate. Currently, RS is limited by its specificity. False positive spectra in RS arise most often from tissues with high nuclear density such as the epidermal layer of skin, hair structures, and areas of inflammation.^9,10^

Reflectance Confocal Microscopy (RCM) is a complementary optical imaging technique that offers noninvasive visualization of skin structures *in vivo* at subcellular resolution,^11^ and it has been advanced into clincial practice for noninvasive guiding diagnosis of skin cancer.^12^ RCM was first reported to be used on BCC by González *et al*. in 2002,^13^ and a meta-study has reported the sensitivity of 92% (range 87-95%) and specificity of 91% (range 84-96%) for BCC diagnosis by human using RCM images.^14^ Automatic segmentation on RCM images of different structures were explored by D’Alonzo *et al*.^15^, and they achieved 0.97 area under the curve (AUC) for classification. Automatic diagnosis of BCC by RCM images was also studied by Gabriele Campanella *et al*., and they achieved around 0.89 AUC.^16^

Althogh systems using the combination of RS and RCM has been developed,^17,18^ the utilization of this combination has not been studied for the diagnosis of BCC. We proposed a method to combine RS and RCM to improve the diagnostic accuracy of BCC diagnosis. Our previous report using Raman spectroscopy alone^19^ discriminated BCC from normal tissues with 100%/84% sensitivity/specificity (30 patients, 223 spectra).^20^ Many of the false positives originate from hair structures, epidermis, and inflammation. Further examination of the RCM images that were acquired at the same site as the RS data showed that hair structures and epidermis are often easily discernable in the structural RCM images. Hair exhibit circular structure from the shaft cross section, and epidermal layer exhibits ribbon patterns from the highly scattering stratum corneum and honeycomb shaped cellular patterns whitin the squamous epithelium. Therefore, we trained two Convolutional Neural Network (CNN) models to identify hair structures and epidermis, respectively. After training, we collected all the RCM images corresponding to positive predicted RS (including all true positive and all false positive) and used the two CNN models to identify hair structures and epidermis images and remove the corresponding sites from the positive to the negative category. By doing so, we converted more than 54% of the false positves to true negatives, thus increasing specificity to 93% (an 8% increase) while keeping the sensitivity at 100%.

## 2 Methods

### 2.1 Clinical data set

For this analysis, we combined RS and RCM images from two prior studies (Feng 2019^20^ and Feng 2020^21^) consisting of 292 site-matched RS and RCM images (141 and 151, respectively for the two papers) from Mohs surgical sections. Fig. 1 illustrates a typical experiment whereby an RS spectrum and RCM image are acquired from the exact same location over various skin constituents including hair structures, dermis, epidermis, fat, inflammation, and basal cell carcinoma (BCC). Raman spectra and RCM images were collected using a custom-built Raman confocal microscope with 830 nm excitation wavelength.^19^

**Fig. 1.**
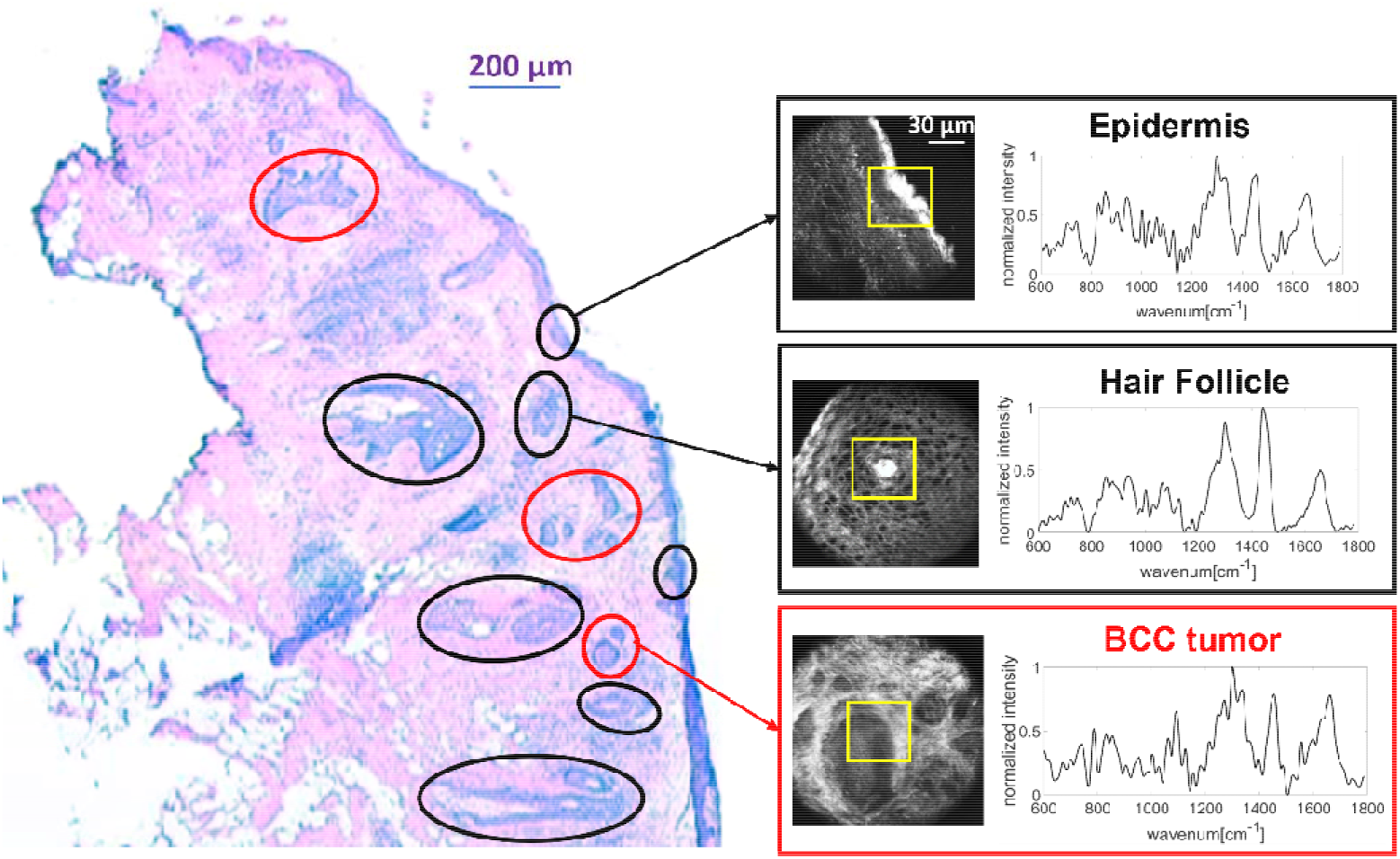
Collection of Raman spectra and RCM images from a BCC tumor section. Red circle: BCC tumor. Black circle: normal tissue (hair structure or epidermis). Yellow square: regions that were assessed by Raman scanning.

The Feng 2019 study reported Raman imaging on 30 frozen tissue blocks from 30 patients. A previously developed biophysical model^19^ was used on Raman spectra to discriminate BCC and normal tissues and achieved 100% sensitivity and 84% specificity when prioritizing sensitivity. RCM imaging and frozen section hematoxylin and eosin (H&E) staining were also acquired for reference, but were not used for diagnosis.

Figure 2 describes the RCM images used for this study. The RCM images in the “task group correspond to the Raman spectra that were predicted as containing BCC, including both false positives and true positives, when prioritizing sensitivity. The rest of the RCM images are in the “train group”. To enlarge the train group, we also used the RCM images from the Feng 2020 study.^21^ Those images were obtained using the same system and in the same scale as the Feng 2019 study. Overall, there were 230 RCM images (30 BCC and 200 non-BCC) in the train group and 62 RCM images (36 BCC and 26 non-BCC) in the task group. Furthermore, we removed 39 images of hair structures (including hair follicle, hair shaft, and hair core) from the train group that had abnormal patterns and looked very different from the other hair structure images, which resulted in 191 RCM images in the final train group. There were three reasons why an image was labeled abnormal. First, the image frame failed to cover the complete circular hair structure (e.g., only the outer arc was visible in the image). Second, large areas of pixels were missing because of problems in the acquisition process. Third, some hair structures had an abnormal growth appearance that did not conform to the regular circular shape (Appendix A presents all images that were noted as abnormal and removed from the train set).

**Fig. 2.**
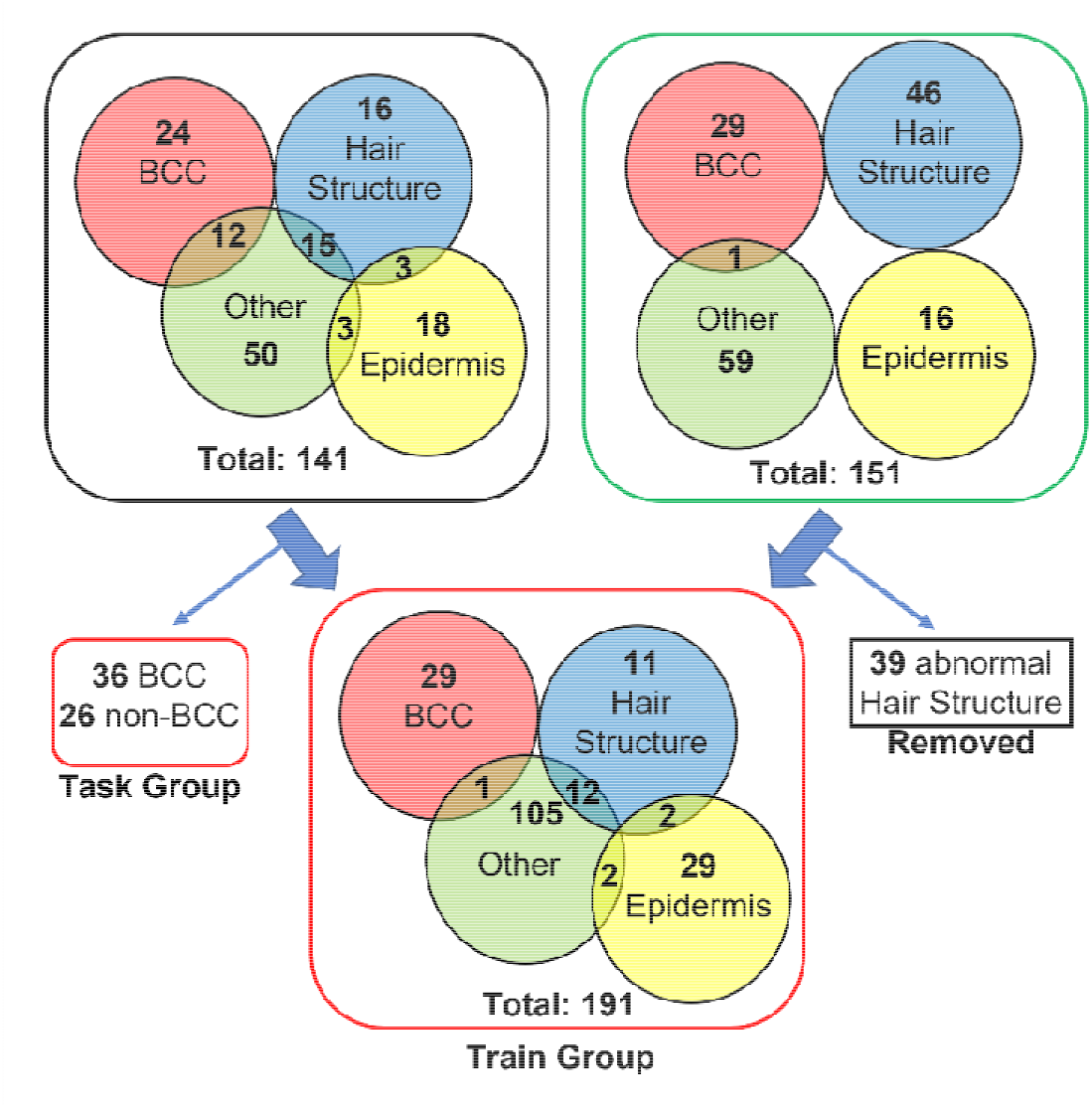
Description of the RCM image sets used for the train and task groups. Images in the round black box were from Feng *et al*.’s Raman biophysical model paper^20^, images in the green box were from Feng *et al*.’s Raman super-pixel paper.^21^ Images in the red boxes were Train group and Task group used for our models. Images in the rectangular black box were the abnormal hair structure images that were removed.

### 2.2 Data Preprocessing

To prepare images for use in training the CNN, we performed several preprocessing operations to the images in the “train group”. First, we randomly divided the images in the “train group” into training set, validation set, and test set. We trained two binary classification models. One model is trained to classify “hair structure” vs. “non-hair structure”; the other one is trained to classify “epidermis” vs. “non-epidermis”. However, both categories are unbalanced in the training set (23 hair structure images and 168 non-hair structure images, 33 epidermis images and 158 non-epidermis images). Therefore, in only the training set, we performed data augmentation on both categories for the two models respectively (when training the hair structure model, we only augmented hair structure images, and when training the epidermis model, we only augmented epidermis images) by flipping, rotating, scaling, and shifting. After positive category augmentation, we resized all images to 224*224*3 (height*width*channel) and performed routine normalization and standardization (each pixel minus mean value and divide by standard deviation). Although these were grayscale images, we kept 3 RGB channels to match the input size for transfer learning.

### 2.3 Model Architecture and Training

Given our relatively small training set, transfer learning is a suitable choice for our models. Transfer learning uses the weights of a pre-trained network to transfer into our own models and adding only a small number of layers with a small number of parameters to be trained on our data.^22^ This approach has been demonstrated as a powerful tool in medical imaging field.^23–26^ We used ResNet50^27^ as the model architecture. We removed the top fully connected layers and added one fully connected (FC) layer with two neurons as output. The activation function was SoftMax. The loss was calculated using cross-entropy.

To train the hair structure model, we imported all weights for the 50 layers trained from ImageNet and left the FC layer trainable. To train the epidermis model, we imported all weights for the first 40 layers trained from ImageNet and left the rest (10 layers) and the FC layer trainable. For both models, we used mini-Batch Stochastic Gradient Descend to minimize the cross-entropy loss and used momentum with /3 = 0.9 as the optimizer. In addition, we added L2 loss with A = 0.01 for regularization. When inputting images, we randomly selected a batch of 16 images and performed augmentations by flipping, rotating, shifting, and scaling. All the augmentations were randomly selected and were applied on random images in the batch. This function was fulfilled by TensorFlow ImageDataGenerator. The trainings were performed with one NVIDIA Tesla K80 Graphic Card provided on Google Cloud Platform. Each model was trained for 0.5 hour with 100 epochs.

The final diagnosis was made based on the union of the two models. Within the “task group”, we chose the images that were classified as hair structure or epidermis by the two models and converted their corresponding spectra from positive to negative.

## 3 Results

### 3.1 Training Process

The model training process is shown in Fig. 3. Hair structures are more complex structures than epidermis, so training the model to identify hair structures required more epochs to converge than the epidermis model did, whose loss started increasing after about 25 epochs because of overfitting. The receiver operating characteristic (ROC) curves on test sets showed good performance of the models (area under the curve of 0.99 for the hair structure model and 0.93 for the epidermis model). We only used positive sample augmentation on the training and validation sets, so the number of positive and negative samples was still unbalanced in the test set, and the total sample number was small. For example, the test set for the hair structure model had only 5 hair structure (positive) samples and 30 non-hair structure (negative) samples. Thus, the number of operating points on the ROC curves for the test sets was limited (Figure 3).

**Fig. 3.**
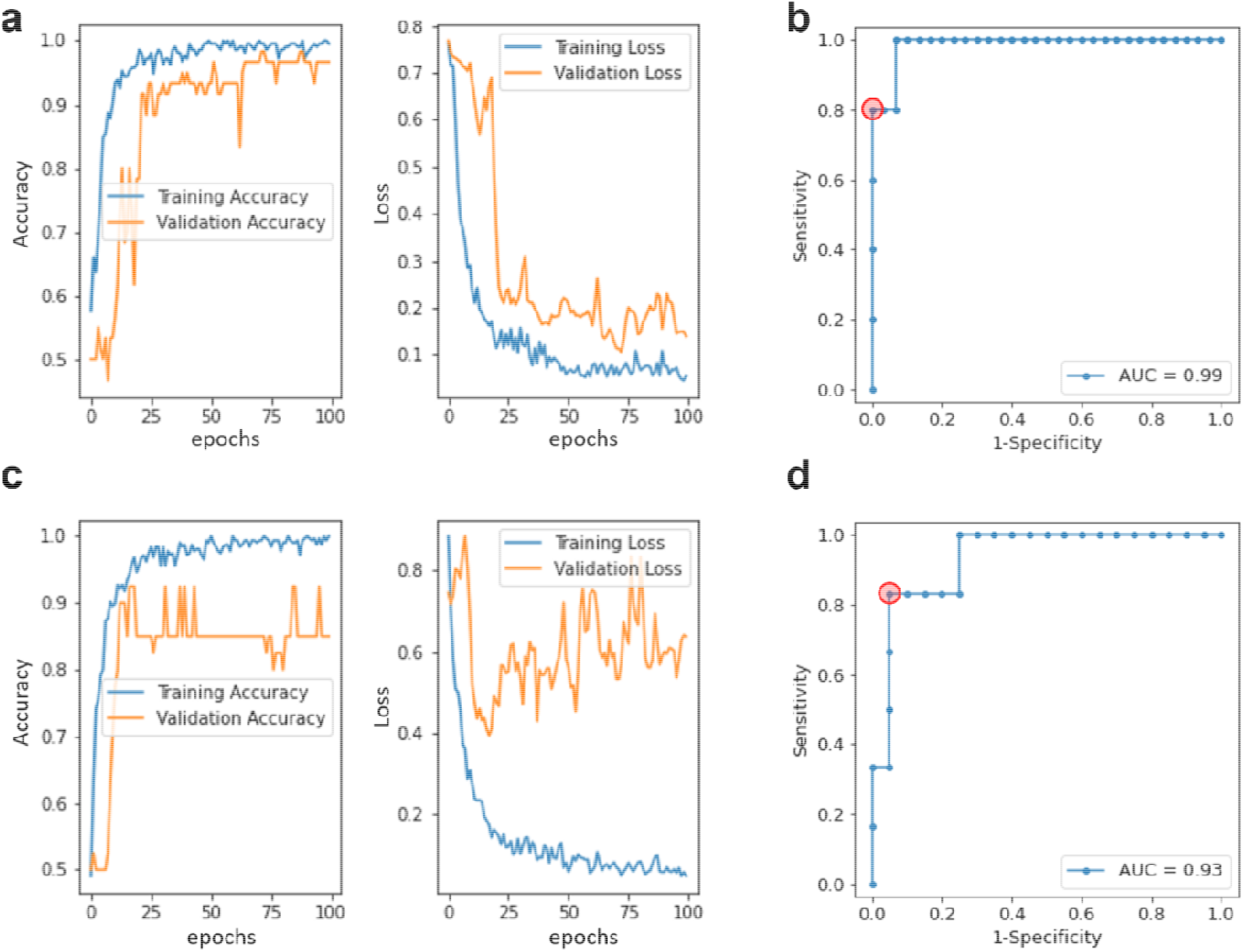
Training process details. (a) Training and validation loss and accuracy for hair structure model. (b) ROC curve for hair structure model on test set. (c) Training and validation loss and accuracy for epidermis model. (d) ROC curve for epidermis model on test set. The red circles in (b) and (d) are the operating points corresponding to the thresholds selected for use in subsequent model validation on the task group data.

### 3.2 Model validation

We applied both models on the independent task group. The thresholds for each model were selected based on the ROC curves on the test set in Fig. 3. We chose thresholds to achieve high specificity and acceptable sensitivity. For the hair structures model, we selected the threshold that provided sensitivity/specificity of 80%/100% on the test set, and for the epidermis model we selected the threshold that provided sensitivity/specificity of 95%/84% on the test set. The hair structure model predicted 8 non-BCC images and 0 BCC images as hair structures for the task group. The epidermis model predicted 5 non-BCC images and 0 BCC images as epidermis for the task group. Overall, there were 13 non-BCC images chosen by the models as non-BCC and 0 images identified as BCC for the task group. Fig. 4 (a) illustrates exemplar images from the task group that the models identified as containing hair structures or epidermis. The circular and the ribbon structure are obvious in these images. Of the 26 false positive spectra from the Raman model (Fig. 4b), the hair structure and epidermis models identified 13 images (associated with 14 spectra) to be either hair structures or epidermis. Therefore, 14 spectra were converted from false positives using Raman alone to true negatives by considering RCM. This reduced the false positives by 54% while keeping the sensitivity at 100% (Fig. 4b).

**Fig. 4.**
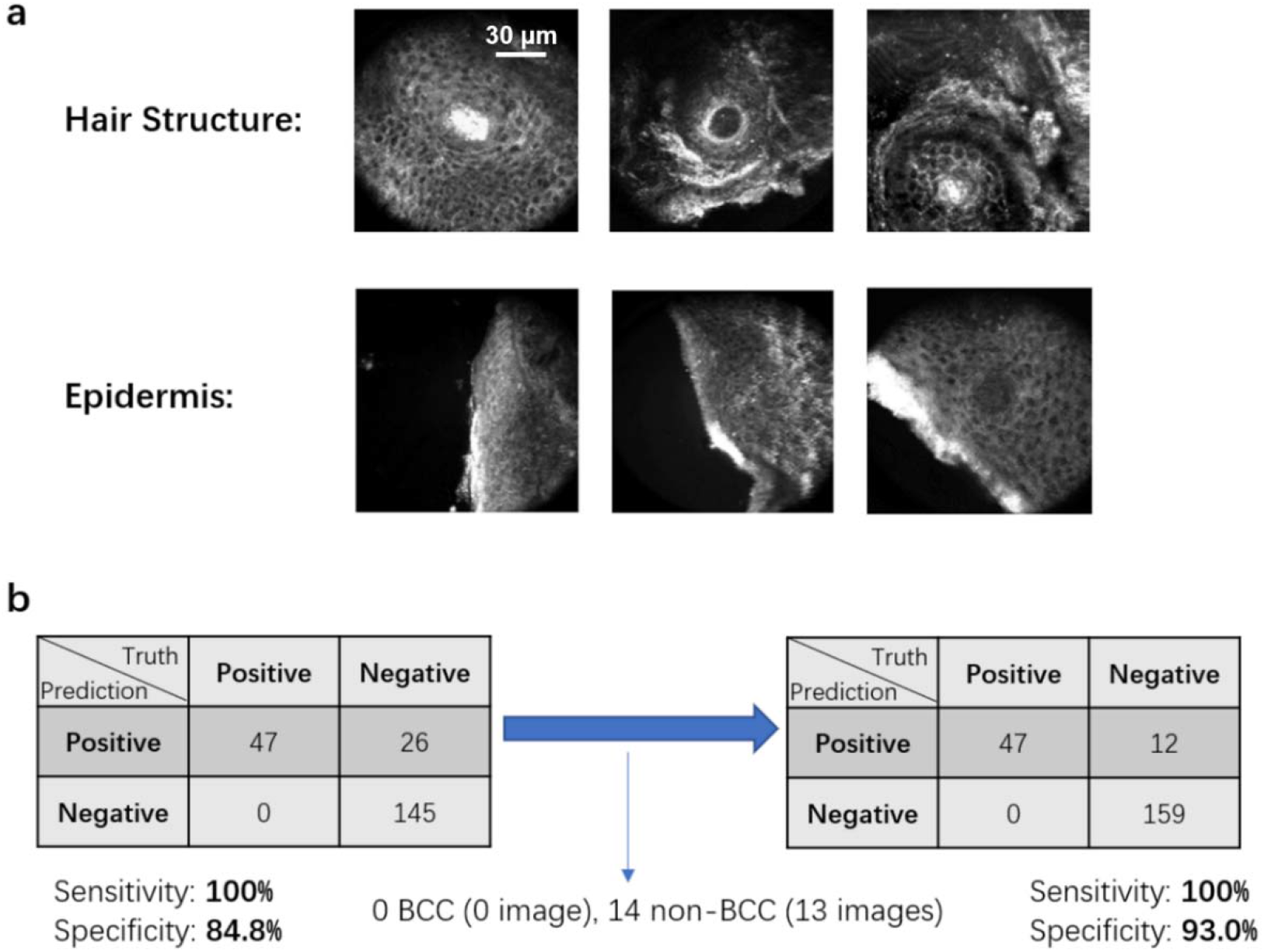
(a) Illustration of the images within task group that were identified as epidermis by the epidermis model and as hair structures by the hair structure model. (b) The confusion matrix for the Raman analysis changes after applying the models to identify epidermis and hair structures.

## 4 Discussion

This study demonstrates the feasibility of combining Raman spectra and RCM images for skin cancer diagnosis. By training two CNN models with transfer learning to recognize hair structure and epidermis on RCM images, we removed 54% of the false positives from Raman while maintaining 100% sensitivity. The RCM models can accurately identify images containing circular patterns of hair structures and ribbon patterns of epidermis.

When preprocessing the training data, we removed hair structure images depicting les common shapes. Although this helped the model to identify the typical circular shapes of most hair structures on RCM images, hair structures that have less common shapes cannot be identified by the model. This can be improved in future studies by expanding the dataset to include a larger number of examples, especially of images depicting less common hair structures.

We used RCM images to identify hair structures and epidermis, not for BCC diagnosis. We chose this approach because the previous Raman biophysical model provided an interpretable prediction on BCC tissues, and the obvious structure of hair and epidermis in RCM images makes our current model reliable and easy to train. The combination of the two models would be both reliable and interpretable. Another approach to combining RS and RCM could use multimodal learning, where RS and RCM images serve as inputs into the single model and for BCC classification.

## Appendix A

Fig. S1 shows all the abnormal hair structure images we removed from our training group.

**Fig. S1.**
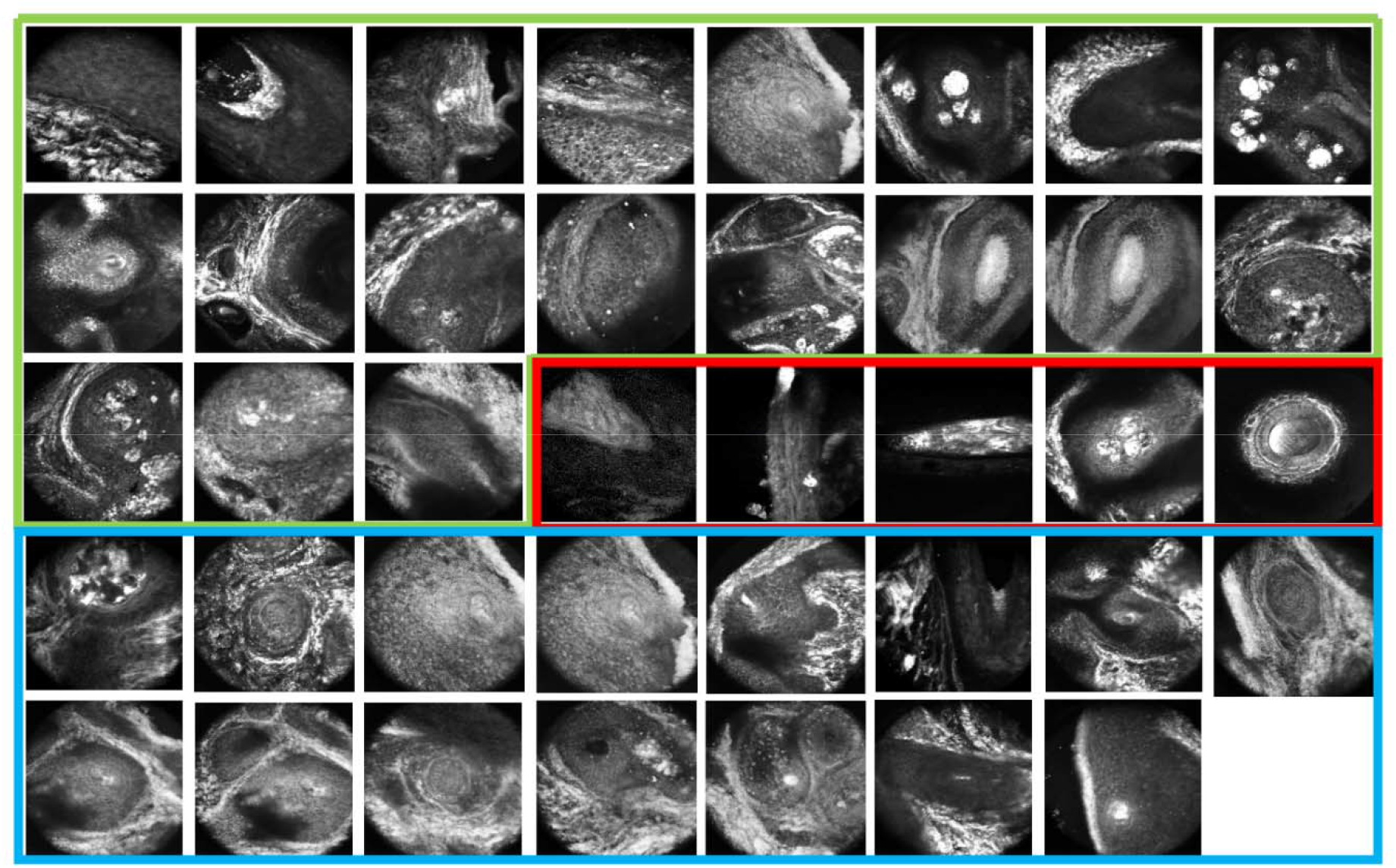
Abnormal hair structure images we removed from our training group. Images in the green box were removed for reason 1: they failed to cover the complete circular hair structure. Images in the red box were removed for reason 2: large areas of pixels were missing because of the problems in the acquisition process. Images in the blue box were removed for reason 3: some of the hair structures were abnormally grown on the tissues.

Fig. S2 show all the false positive images and those were selected by the models. In the red box are the images selected by the models and in the blue box are those not being selected. The figure demonstrates we have selected almost all images with ribbon structure (epidermis) and circular structure (hair).

**Fig. S2.**
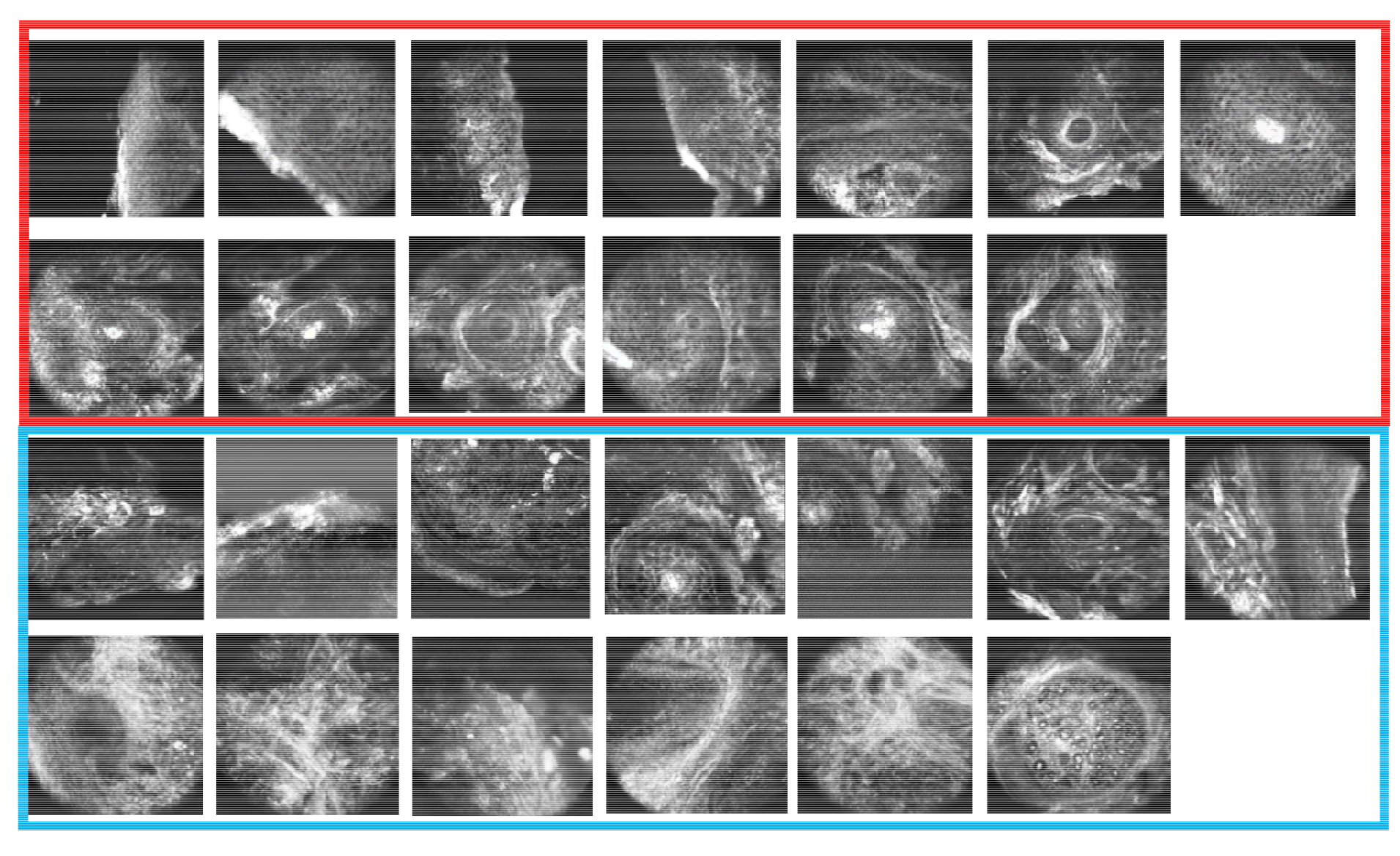
All the false positive images. Images in the red box were selected by the models and removed form false positives. Images in the blue box were not selected by the model.

